# An accurate assignment test for extremely low-coverage whole-genome sequence data

**DOI:** 10.1101/2021.06.04.447098

**Authors:** Giada Ferrari, Lane M. Atmore, Sissel Jentoft, Kjetill S. Jakobsen, Daniel Makowiecki, James H. Barrett, Bastiaan Star

**Affiliations:** Centre for Ecological and Evolutionary Synthesis, Department of Biosciences, University of Oslo, Blindernveien 31, NO-0371 Oslo, Norway; Department of Environmental Archaeology and Human Paleoecology, Institute of Archaeology, Nicolaus Copernicus University, Poland; McDonald Institute for Archaeological Research, Department of Archaeology, University of Cambridge, UK; Department of Archaeology and Cultural History, NTNU University Museum, Trondheim, Norway

**Keywords:** Population assignment, genome skimming, haplotype, chromosomal inversion, ecotype

## Abstract

Genomic assignment tests can provide important diagnostic biological characteristics, such as population of origin or ecotype. In ancient DNA research, such characters can provide further information on population continuity, evolution, climate change, species migration, or trade, depending on archaeological context. Yet, assignment tests often rely on moderate- to high-coverage sequence data, which can be difficult to obtain for many ancient specimens and in ecological studies, which often use sequencing techniques such as ddRAD to bypass the need for costly whole-genome sequencing. We have developed a novel approach that efficiently assigns biologically relevant information (such as population identity or structural variants) in extremely low-coverage sequence data. First, we generate databases from existing reference data using a subset of diagnostic Single Nucleotide Polymorphisms (SNPs) associated with a biological characteristic. Low coverage alignment files from ancient specimens are subsequently compared to these databases to ascertain allelic state yielding a joint probability for each association. To assess the efficacy of this approach, we assigned inversion haplotypes and population identity in several species including Heliconius butterflies, Atlantic herring, and Atlantic cod. We used both modern and ancient specimens, including the first whole-genome sequence data recovered from ancient herring bones. The method accurately assigns biological characteristics, including population membership, using extremely low-coverage (e.g. 0.0001x fold) based on genome-wide SNPs. This approach will therefore increase the number of ancient samples in ecological and bioarchaeological research for which relevant biological information can be obtained.

## Introduction

Despite advances in methodologies that allow for the recovery of higher yields of endogenous ancient DNA (aDNA) (e.g., Boessenkool et al., 2017; Carpenter et al., 2013; Gamba et al., 2014; Pinhasi et al., 2015), DNA preservation in sub-fossil, archaeological, historical, or degraded biological material remains variable and is often context specific (Ferrari et al., 2021; Keighley et al., 2021; Tin et al., 2014). In order to account for such unpredictability, aDNA sequencing projects typically screen many specimens from which a subset with the best DNA preservation is selected for deeper sequencing (e.g. Martínez-García et al., 2021; Star et al., 2018; van der Valk et al., 2021). Similarly, DNA (organelle) reference databases are increasingly being generated through “genome skimming” sequencing strategies (e.g. Dodsworth, 2015; Marcus, 2021; Nevill et al., 2020; Zeng et al., 2018). These practices result in a proliferation of specimens for which (extremely) sparse genome-wide data is obtained (Bohmann et al., 2020). Due to their low coverage, these data are difficult to jointly analyze with specimens that have obtained higher coverage without introducing various types of statistical bias, particularly when dealing with heterochronous datasets (e.g. François & Jay, 2020; Lee et al., 2010; Patterson et al., 2006; Skoglund et al., 2014). For this reason, specimens with low coverage genome-wide data are typically discarded from further bioinformatic analyses, leading to the destruction of unique zooarchaeological specimens for which no meaningful information is obtained. Efforts to obtain as much relevant information as possible from such specimens are therefore important from a biological and an ethical perspective (Pálsdóttir et al., 2019).

Although high-coverage whole genome data allow a plethora of detailed bioinformatic analyses, high coverage is not necessarily required for the determination of basic biological characteristics. Moreover, depending on archaeological context, knowledge of such characteristics can provide highly relevant information on population continuity, species migration and distributions, hunting, and historic trade and/or burial practices. For instance, the genetic sex of ancient mammals can easily be determined from sparse sequencing data due to its association with extensive genomic differentiation on a chromosomal scale. Sexing has been applied to ancient low-coverage sequences to infer burial practices (Fages et al., 2020; Nistelberger et al., 2019), the impact of historic hunting (Barrett et al., 2020), and the behaviour of extinct species (Pečnerová et al., 2017).

Other relevant biological characteristics may also be associated with large scale genomic differentiation. In particular, structural variants (e.g. chromosomal inversions) have been increasingly identified as major drivers of evolutionary and ecological processes (Wellenreuther & Bernatchez, 2018), playing important roles in population structure and evolution. For instance, inversions are involved in the evolution of sex chromosomes (Hughes et al., 2010; Lemaitre et al., 2009) and speciation (Noor et al., 2001), and are critical for within-species adaptation to local environments (Ayala et al., 2013; Barth et al., 2017; Jones et al., 2012; Leitwein et al., 2017; Lowry & Willis, 2010; Morales et al., 2019; Nadeau et al., 2016; Pettersson et al., 2019; Todesco et al., 2020; Twyford & Friedman, 2015). Chromosomal inversions can affect megabase sized genomic regions (e.g. P. R. Berg et al., 2017; Fang et al., 2012; Twyford & Friedman, 2015), and are often characterized by high levels of linkage disequilibrium (LD) (Hoffmann & Rieseberg, 2008) due to inhibited recombination between non-collinear inversion haplotypes. Thus, genotyping of such haplotypes using a subset of segregating genetic markers is feasible using whole genome sequencing data (Donnelly et al., 2010; Salm et al., 2012).

Several methods have been developed for assigning inversion haplotypes in order to facilitate GWAS analysis for SNPs within inversions in the human genome (*scoreInvHap*, Ruiz-Arenas et al., 2019; *pfido* Salm et al., 2012; *InvClust*, Cáceres & González, 2015; *inveRsion*, Cáceres et al., 2012). These methods rely on LD break-points and structural variation (e.g. *InvClust, inveRsion*, and *scoreInvHap*, as well as methods proposed by (Bansal et al., 2007; Sindi & Raphael, 2010) or haplotype tagging (*inveRsion*) to identify inversion sites and then conduct various types of SNP calling within those sites. All of these methods are specifically developed for identifying inversions in human genomes (e.g., Ma & Amos, 2012) and their use in disease- and other phenotype-association studies (Ruiz-Arenas et al., 2019; Salm et al., 2012). They have not been tested with sparse genomic data and are specific to use with inversions; indeed, *pfido* was designed for just one inversion in the human genome (Salm et al., 2012). Because of their reliance on signatures of structural variation, they cannot be applied to other types of variation, such as genome-wide population differentiation. There is currently no approach specifically designed to classify extremely low-coverage data with a broad applicability to score different types of large-scale genomic differentiation in a range of species.

Here, we developed a new method that allows efficient assignment of different biological characteristics using extremely low-coverage sequence data. The approach is similar to *scoreInvHap (Ruiz-Arenas et al., 2019)* in that scoring is a two-step process, yet there are some key differences. First, a database is created with the allele frequency association of individual SNP loci with a specific biological character (e.g. an inversion type or population membership). These databases are based on moderate- to high-coverage sequences of a subset of specimens (Figure 1a). Second, sequence alignment data of (ancient) specimens are compared to this database and a joint probability (e.g. see Star et al., 2017) is calculated based on the binomial distribution of their frequency association (Figure 1b). Importantly, in contrast to earlier approaches, this probability calculation does not make any assumptions regarding specific signatures of structural variation and can therefore be applied to different types of genetic differentiation. This includes differentiation between inversion haplotypes or genome-wide differences associated with ecotype or population structure. Our program depends solely on freely available, commonly used software and file formats and is freely available for download at: https://github.com/laneatmore/BAMscorer.

**Figure 1.**
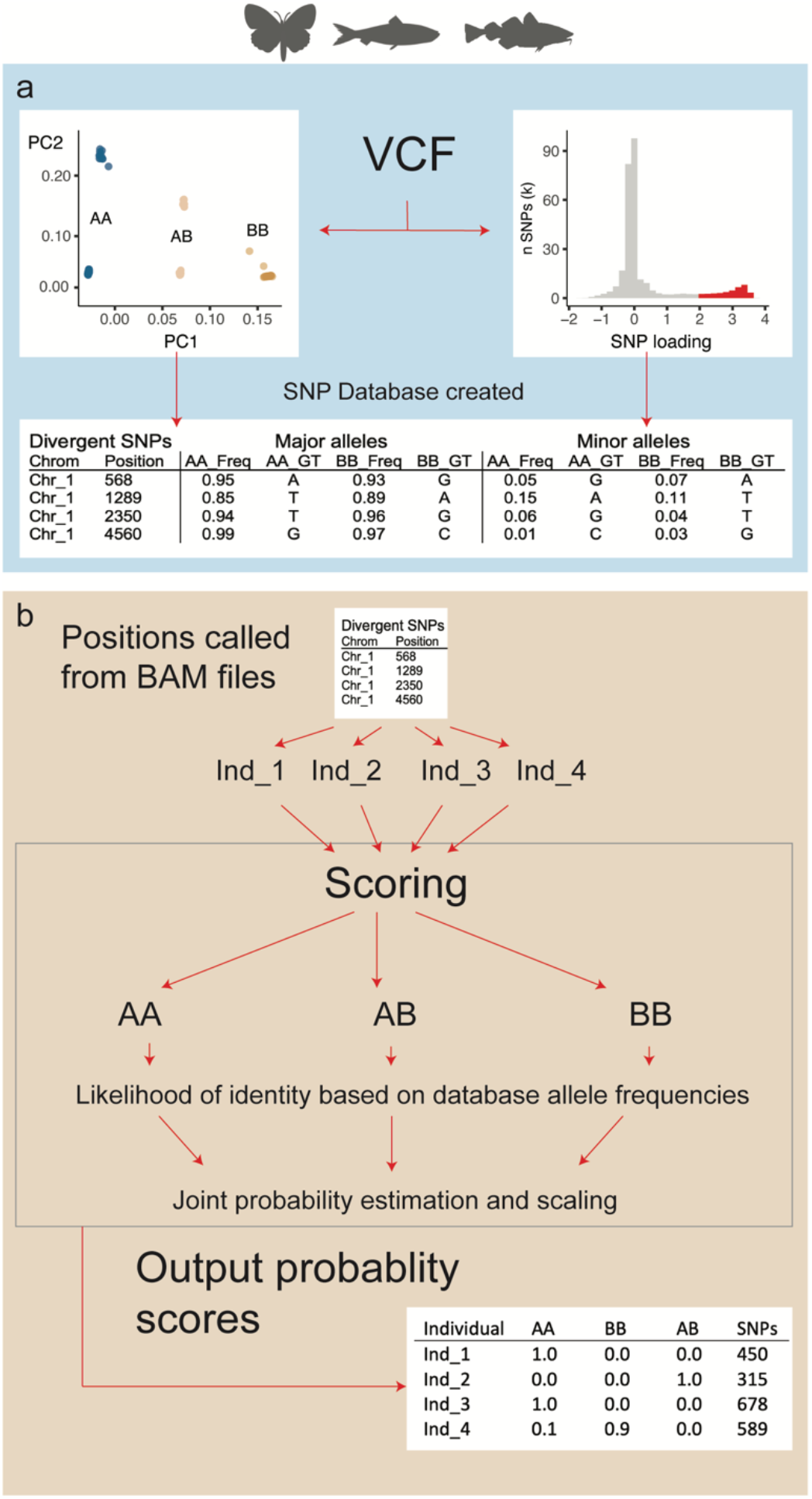
The BAMscorer pipeline. The BAMscorer pipeline has two main modules -- reference database creation and alignment file scoring. a) sequence data must be pre-processed and input into the pipeline as a VCF file. smartPCA is used to generate eigenvalues and SNP loading weights, which are then used to assign haplotypes in the reference dataset and create a database of highly-divergent loci in a given region of interest. b) These positions are called from the alignment files to be scored. The positions are then compared to the database for allelic similarity. The likelihood of a given allele at a locus belonging to a haplotype is coded as the frequency of that allele at the locus in each database. AB allele frequencies are calculated as the average of frequencies present in AA and BB haplotypes. A joint probability is estimated for each alignment file belonging to each of the three haplotypes (for genome-wide assignment only AA and BB are used) and these values are scaled to one, outputting a probability index of genomic assignment for each individual.

We investigated the efficiency of our approach in assigning haplotypes for three chromosomal inversions in species that differ in their availability of reference specimens (P3, *Heliconius numata, n* = 20; Chr12, *Clupea harengus, n* = 19, and LG01, *Gadus morhua, n* = 276). These inversions display clinal distributions that are associated with biological characters such as wing-pattern phenotypes (Joron et al., 2006, 2011; Nadeau, 2016), adaptation to water temperature and salinity (Pettersson et al., 2019) and migratory behaviour (Paul R. Berg et al., 2016). Finally, we investigated the accuracy of this approach for the genome-wide population assignment of western and eastern Atlantic cod specimens (Barth et al., 2019; Pinsky et al., 2021).

## Materials and methods

### Ancient DNA extraction and sequencing

Nine Atlantic herring bones from two sites, dated between the 9th and 15th century CE (Table S1), were UV-treated for 10 minutes per side and cleaned with ultra-pure water. DNA was extracted including a pre-digestion step, following Damgaard et al. (2015). 10-40 mg of bone were pulverized with micro pestles in the digestion buffer (1 ml 0.5 M EDTA, 0.5 mg/ml proteinase K and 0.5% N-Laurylsarcosine). Following overnight digestion, DNA was extracted with 9 volumes of a 3:2 mixture of QG buffer (QIAGEN) and isopropanol. MinElute purification was carried out using the QIAvac 24 Plus vacuum manifold system (QIAGEN) in a final elution volume of 65 μl. Parallel non-template controls were included. Single-indexed blunt-end sequencing libraries were built from 16 μl of DNA extract or non-template extraction blank, following the single-tube (BEST) protocol (Carøe et al., 2018) with the modifications described in Mak et al. (2017). All laboratory protocols up to indexing of sequencing libraries were carried out in a dedicated aDNA clean laboratory at the University of Oslo following standard anti-contamination and authentication protocols (Cooper & Poinar, 2000; Gilbert et al., 2005; Llamas et al., 2017). Library quality and concentration were inspected with a High Sensitivity DNA Assay on the Bioanalyzer 2100 (Agilent) and sequenced on an Illumina HiSeq 2500 platform at the Norwegian Sequencing Centre with paired-end 125 bp reads, demultiplexed allowing zero mismatches in the index tag.

### Data processing

For each species investigated, two different datasets were used. The first dataset was used to create the reference SNPs database and the second dataset, containing different individuals, was scored utilizing the BAMscorer program. All the datasets used in this manuscript —including the newly generated archaeological Atlantic herring data— are publicly available.

#### Heliconius butterflies

The heliconius reference database was created using a set of 20 individuals from various *H. numata* subspecies described in Nadeau et al. (2016). A set of 40 unrelated individuals to be scored were obtained from Jay et al. (2019). Both datasets were aligned to the Hmel2.5 reference assembly (http://ensembl.lepbase.org) using PALEOMIX v.1.2.13 (Schubert et al., 2014) with BWA-mem. Genotypes for the reference database were called using the GATK4 pipeline (Van der Auwera & O’Connor, 2020) and the following filtering parameters: FS<60.0 && SOR<4 && MQ>30.0 && QD > && INFO/DP<5500, SnpGap 10, minGQ 15 minDP 3, maf 0.001, with indels removed and biallelic variants selected. The P3 inversion at the supergene *P* mimicry locus, located on chromosome 15 and associated with wing pattern types in *Heliconius numata* subspecies (Jay et al., 2018; Joron et al., 2011), was investigated. The ~1.1Mb P3 inversion is found on scaffold Hmel215003o (between 2000001-3100000bp).

#### Atlantic herring

The Atlantic herring reference database was created using a set of 21 individuals described in Han et al. (2020), representing all but one of the major herring populations in the eastern Atlantic. The nine ancient Atlantic herring specimens dating from the 9th-15th century in Poland (Domagała & Franczuk, 1992; Iwaszkiewicz, 1991; Makowiecki, 2003; Makowiecki et al., 2016) were scored. Modern herring reads were aligned to the Atlantic herring reference genome (GCA_900700415.1) (Pettersson et al., 2019) as above. Ancient herring reads were aligned as described in Ferrari et al. (2021), using BWA-backtrack. Genotypes for the reference database were called and filtered as described above. Two individual outliers were observed and checked for relatedness using KING (Manichaikul et al., 2010, Supplementary Note 1). These individuals appeared to be duplicates and were removed from the dataset. An ~8Mb inversion on chromosome 12 was investigated, which is associated with different Atlantic herring ecotypes (Pettersson et al., 2019). The inversion is located at chr12:17900000-25600000bp. Ancient herring alignment files were downsampled to 100K reads. Most specimens have excellent DNA preservation (Supplementary Table 1) and all show the typical aDNA fragmentation and misincorporation patterns of authentic ancient DNA data (Supplementary Figure 1).

#### Atlantic cod

The Atlantic cod reference dataset was created using 276 Atlantic cod individuals representing most major geographical locations (western Atlantic, eastern Atlantic, and Baltic sea) in the species’ range (Barth et al., 2019; Pinsky et al., 2021). A dataset of 15 unrelated archaeological specimens were obtained from Star et al. (2017). Modern and ancient reads were aligned to the gadMor2 reference genome (Star et al., 2011; Tørresen et al., 2017) as above. Genotype calling and filtering for the reference was performed as described in Barth et al. (2019) using the GATK haplotype caller v.3.4.46 (McKenna et al., 2010), bcftools v.1.3 (Li, 2011), VCFtools v.0.1.14 (Danecek et al., 2011). An ~16Mb double chromosomal inversion on LG01 which is associated with differences in migratory behaviour (Paul R. Berg et al., 2016; Kirubakaran et al., 2016; Sodeland et al., 2016) was investigated. This inversion is located at LG01:9100000-26200000. Finally, genome-wide data separating 24 western Atlantic from 252 eastern Atlantic cod specimens was analysed excluding the location of four major inversions (on LG01, LG02, LG07, and LG12) following Star et al. (2017).

### Analyses

The BAMscorer pipeline operates as follows:

#### Module 1: Creation of SNP reference databases (Figure 1a)

The initial step of the BAMscorer pipeline is to create a reference database of divergent SNPs associated with each haplotype or population in a set of focal individuals. These divergent SNPs are referred to as “AA” and “BB” haplotypes/groups. SNP databases are created as follows:

1. The VCF file is first prepared with VCFtools v.0.1.16 (Danecek et al., 2011) and PLINK v.1.9 (Purcell et al., 2007), selecting only those regions of interest (i.e. where inversions are located or genome-wide).
2. A Principal Component Analysis (PCA) is run as implemented in smartPCA (EIGENSOFT v.7.2.1, Patterson et al., 2006; Price et al., 2006) to calculate axes of differentiation and individual SNP loadings between inversion haplotypes or populations. As a default, the BAMscorer pipeline selects diagnostic loci in the top and bottom 5% of the SNP loading distribution, although the optimal SNP loading cut-off value should be determined by the user. Visualization of the SNP-loading profile can help decide such cut-offs (see further below).
3. SNPs that pass cut-off filters form the divergent SNPs database for each haplotype or population. To help make relevant selection of individuals, heterozygosity is calculated per individual based on SNPs in the divergent database.
4. Individuals from the VCF file are scored for PC1 and heterozygosity values and manually classified into types: homozygous haplotypes AA and BB, and, if applicable — i.e. in the case of inversions —, heterozygous AB. Inversion haplotypes are known to fall into specific clusters in PCA analysis (see Figure 1a), which allow for easy identification using separation on PC1 and assessing levels of individual heterozygosity.
5. For individuals in AA and BB haplotypes, allele frequencies of the divergent SNPs are calculated. Two databases are created, containing the allelic state (e.g. A, C, G, T) and allele frequencies of the major (first database) and minor (second database) alleles in the AA and BB haplotypes. Databases containing few individuals can contain fixed alleles due to limited sampling. This uncertainty in sampling fixed alleles is addressed by calculating an expected frequency of (1/((2*N)+ 1)) where N is the number of individuals in the reference database for fixed alleles in the region of interest. When scoring inversions, allele frequencies for AB haplotypes are averaged to approximate the probability of observing a random set of alleles coming from either AA or BB haplotype

Once optimal database parameters have been identified (a full list of parameters can be found at https://github.com/laneatmore/BAMscorer), the SNP database can be reused for BAM scoring on many different datasets of the same species.

#### Module 2: BAM scoring (Figure 1b)

1. The divergent SNPs databases are used to score alignment files (BAM format) for a given set of (low-coverage) individuals. For each locus in the divergent SNPs database, matching reads are pulled from the BAM file using the python module pysam (https://github.com/pysam-developers/pysam). Allelic state is determined based on the most highly-represented allele in all reads for each position. In the event that there are equal numbers of reads for multiple alleles at a given locus, one allele is then chosen at random. This process provides a subset of observed alleles at divergent loci in each inversion or population for each individual BAM file.
2. The probability of observing an SNP variant associated with inversion haplotype or populations is based on allele frequencies of matching positions in the reference databases, e.g. if the position in the BAM file matches the dominant allele in haplotype AA, the probability for that locus in the BAM file is coded as the allele frequency of the dominant allele in haplotype AA. For each position, three probabilities are recorded -- the frequency of that allele in haplotypes AA, BB, and AB (only AA and BB for genome-wide analysis).
3. Joint probabilities of all observed alleles belonging to a particular haplotype or population are calculated for each individual using the following equation:

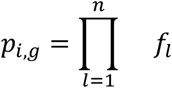

Whereby the probability (*p*) of the scored individual (*i*) and genotype (*g*) is the product of allele frequencies (*f*) of the number (*n*) of observed SNP loci (*l*) in each database.
4. Finally, the joint probability scores for all genotypes are scaled to one to provide a final probability estimate of an individual belonging to a certain haplotype or population. We also report the number of SNPs in the reference dataset that were recovered from each individual BAM file to provide information on how well scored a specific individual is.

#### Assessing scoring certainty

To investigate the sensitivity of the BAMscorer pipeline, we downsampled each BAM file in the three datasets (Heliconius, Jay et al. 2019, Atlantic herring and Atlantic cod, Star et al. 2017). Following an approach described in Nistelberger et al. (2019), BAM files containing whole genome shotgun data were downsampled to contain between 1K and 40K reads (in most instances this is a mere fraction of the available data). At each read interval, and for each individual, the down-sampling was randomly iterated 20 times. We compared accuracy of the scoring results of the extremely down-sampled Heliconius data by comparing results to an independent PCA analysis of the complete dataset (Supplementary Figure 2). For Atlantic herring and Atlantic cod accuracy of results was confirmed by prior knowledge of the inversion haplotypes or geographic origin of specimens.

## Results

We investigated three chromosomal inversions and one genome-wide analysis using BAMscorer. The Heliconius P3 inversion is the smallest (1.1Mb) inversion, followed by the Atlantic herring Chr12 (8Mb) and Atlantic cod LG01 inversion (16Mb, Table 1). Principal Component Analysis (PCA) as implemented through BAMscorer *-select_snps* separates the three main inversion genotypes along PC1 for the Heliconius P3, Atlantic herring Chr12 and Atlantic cod LG01 datasets (Figure 2a), reproducing earlier observations (Barth et al., 2019; Han et al., 2020; Nadeau et al., 2016; Pinsky et al., 2021). Similarly, the whole genome analysis separates western from eastern Atlantic cod specimens along PC1 (Figure 2a, Pinksy et al. 2021). For the data analysed here, BAMscorer *-select_snps* typically runs within 15 minutes. The SNP weight loading distribution underlying genetic divergence between inversion haplotypes of populations is either approximately symmetrical (e.g. Heliconius or Atlantic herring) or asymmetrical (Atlantic cod, Figure 2b). SNP weights are proportional to the correlation (across samples) between each SNP and each PC (Patterson et al., 2006; Price et al., 2006). SNPs that are strongly associated with divergence will have the highest SNP weight loading values and are therefore biologically informative.

**Table 1.** Inversion and genome characteristics of Heliconius, Atlantic herring and Atlantic cod. Each comparison differs in terms of size of inversion, overall genome size and relative size of inversion in terms of species-specific genome size, as well as in terms of the optimum number of divergent SNPs (see methods) and individuals used for the reference databases and scoring.

| Species | Inversion name, location | Inversion size (Mbp) | Genome size (Mbp) | Relative size (%) | Divergent SNPs (n) | Database individuals (n) | Scored individuals (n) |
| --- | --- | --- | --- | --- | --- | --- | --- |
| *H. numata* | P3, Chr15 | 1.1 | 273 | 0.4 | 1701 | 20 | 40 |
| *C. harengus* | Chr12 | 8 | 726 | 1.1 | 28205 | 19 | 9 |
| *G. morhua* | LG01 | 16 | 643 | 2.5 | 47564 | 276 | 15 |
| *G. morhua* | WG | na | 643 | na | 221790 | 276 | 15 |

**Figure 2.**
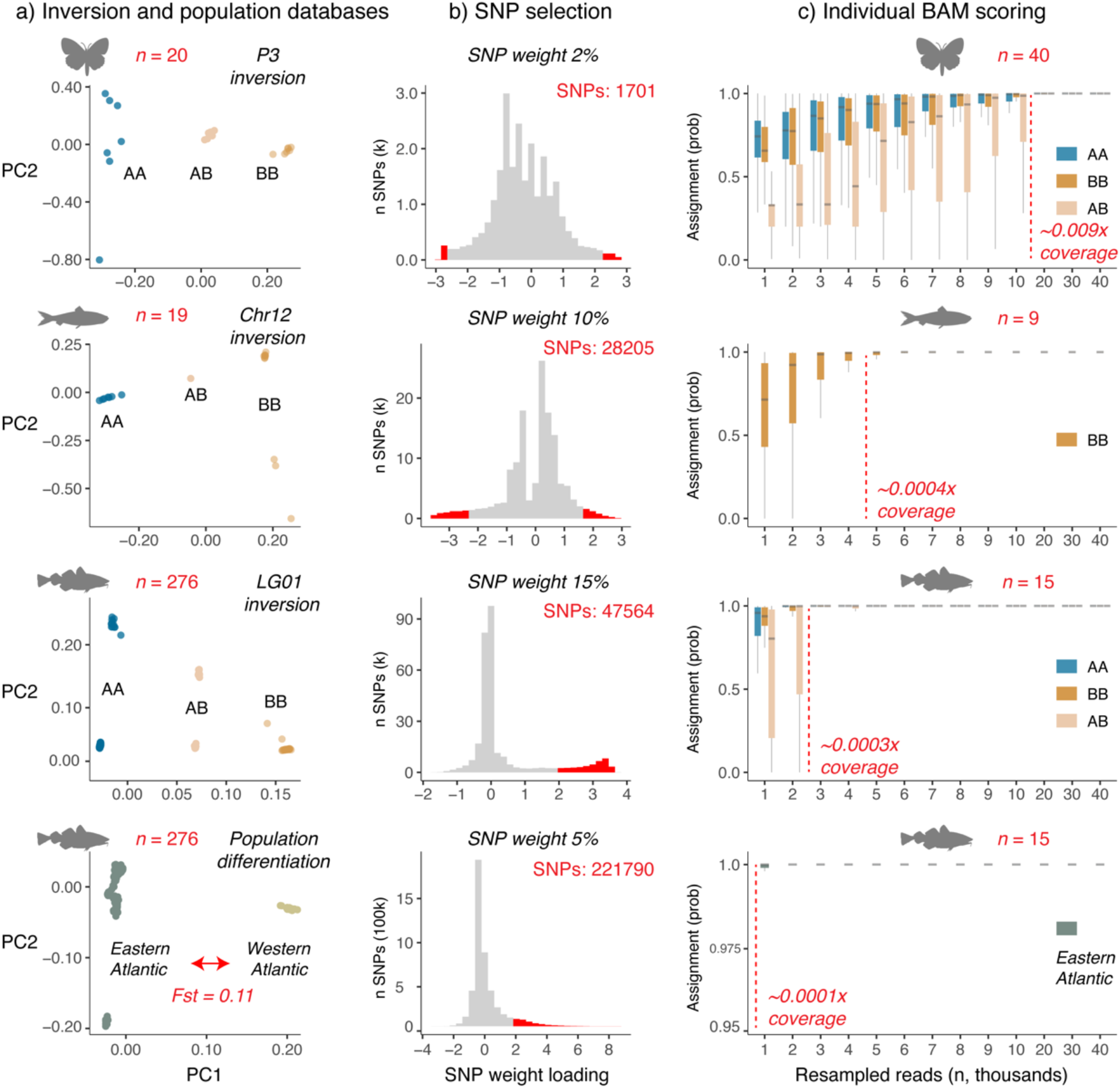
Inversion and population assignment for *H. numata* (P3 inversion), *C. harengus* (Chr12 inversion) and *G. morhua* (LG01 inversion, population differentiation) using extreme low coverage data. a) Inversion and population PCA plots generated for the three species (silhouettes) using smartPCA (ref). The number of individuals (red) and the genome-wide *Fst* differentiation for *G. morhua* (red arrow) is indicated. b) SNPs most associated with either inversion (A or B) haplotype or large-scale population differentiation (western or eastern Atlantic) are selected based on their SNP weight loading distribution along PC1. Those with lowest and highest loadings are most associated with differentiation along PC1. SNP weight indicates the percentage of SNPs selected from the most extreme end(s) of the distribution (red). c) Assignment probability for individual specimens generated by down-sampling BAM files 1000 to 40 000 reads. At each interval, and for each individual, the down-sampling is iterated 20 times in order to generate box plots. Probabilities are calculated based on the joint binomial distribution of observing divergent SNPs associated with either genotype or population. Also indicated is the number of individuals scored (red, note these are not the same individuals used to create the original databases) and fold coverage (red dotted line, x coverage) at which more than 0.99 median assignment probability is obtained.

An important consideration of our approach therefore lies in the selection of loci based on their SNP-loading distribution patterns. In order to maximize the probability of observing loci in low-coverage sequencing data, as many loci as possible should be included in the database. Yet adding those loci that are not significantly associated with either inversion haplotype or specific population will add noise and uncertainty. We therefore tested the accuracy of our approach using a range of SNP-loading filtering parameters. For inversions, databases were created using cut-off values between 1 and 25%, depending on the species under investigation. For our genome-wide analyses, we set the SNP-loading cut-off weights between 1 and 5%. The default parameter in the BAMscorer pipeline is to take symmetrical portions from each side of the SNP-loading distribution (the 5% cut-off value, therefore, takes the top *and* bottom 5% of SNPs), yet we also noticed asymmetrical SNP-loading distribution values. We therefore also investigated the effect of selecting SNPs from either the top or bottom of the SNP-loading distribution.

For instance, for the Heliconius P3 inversion, the ability to confidently score heterozygous individuals (Jay et al. 2019) erodes with increasing SNP weight values (Figure 3), and the optimal cut-off to simultaneously score all possible genotypes lies at 2% and 1701 SNPs. For Atlantic herring Chr12, not all haplotypes are observed in the ancient read data, yet no major increase in ability of scoring is obtained after a SNP weight of 10% and 28205 SNPs (Supplementary Figure 3). For Atlantic cod, best separation of ancient data (Star et al., 2017) was obtained by selecting SNPs from the single, most extreme end of the SNP weight loading distribution (Figure 2B, Supplementary Figures 4-6). For Atlantic cod LG01, SNP selection is similar to the Heliconius P3 in that the optimal cut-off is a trade-off in scoring homozygotes and heterozygotes, which for cod lies at 15% and 47564 SNPs (Supplementary Figure 5). Finally, best population separation for Atlantic cod is obtained at 5% and 221790 SNPs (Supplementary Figure 6).

**Figure 3.**
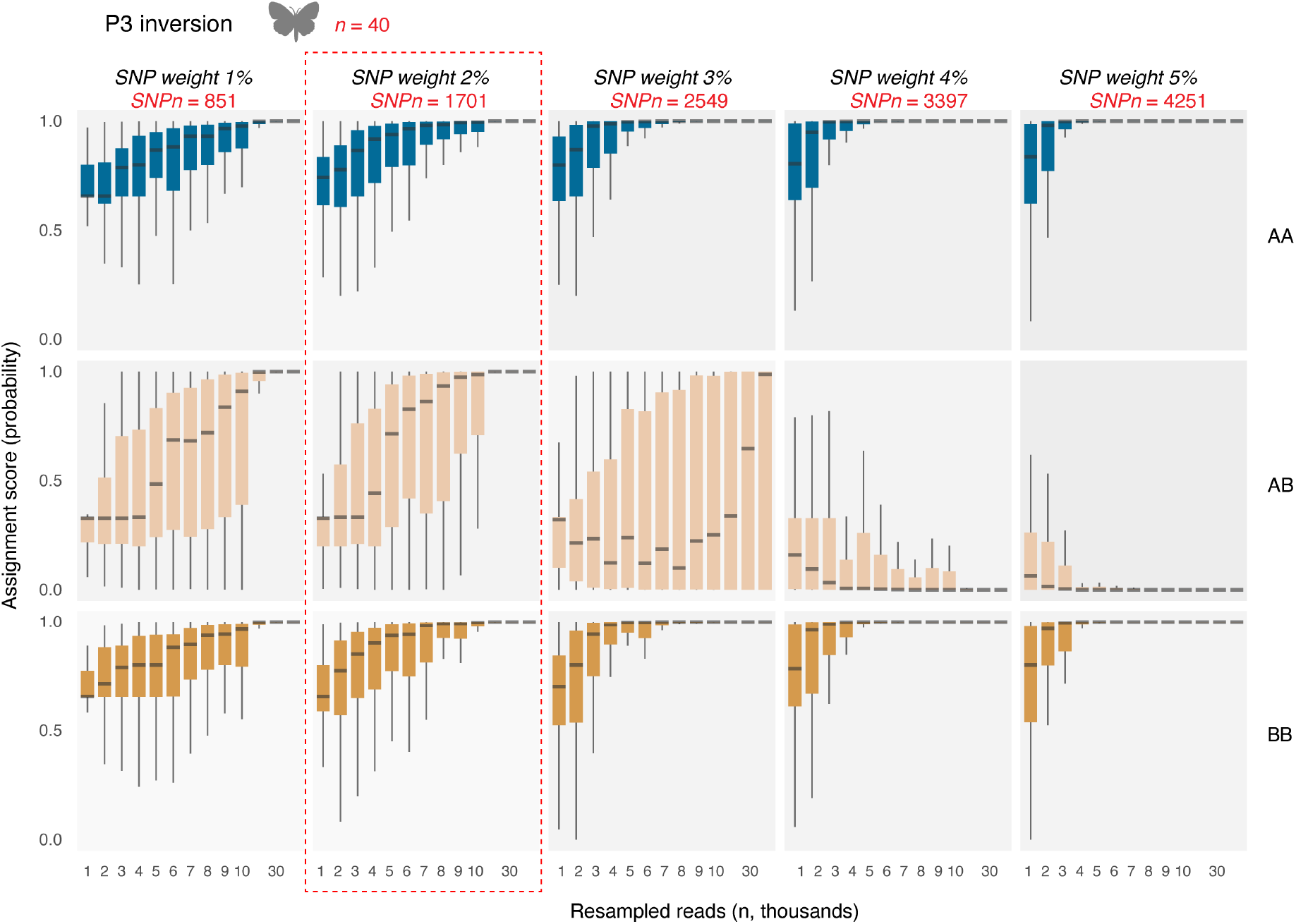
SNP selection by varying SNP weight in *H. numata* (P3 inversion). SNP weight is here defined as the percentage of SNPs with the most extreme values at both sides of the SNP loading distribution. Confidence in probability assignment is obtained by down-sampling BAM files 1000 to 40 000 reads. At each interval, and for each individual, the down-sampling is iterated 20 times in order to generate box plots. Probabilities are calculated based on the joint binomial distribution of observing divergent SNPs associated with either genotype. Also indicated is the number of individuals (n, red) and number of SNPs (SNPn, red) and the chosen cut-off value (red dotted lines) at which all three genotypes can be efficiently separated.

After deciding the best-possible cut-off values, several observations can be made regarding the scoring accuracy of BAMscorer *-score_bams* depending on the number of reads for each of the comparisons. First, accurate scoring is obtained in extremely low coverage data for all comparisons (Figure 2c). For Heliconius, accurate genotype determination was obtained with 20K reads and 0.009x fold nuclear coverage. For all other comparisons, even less reads *—*by an order of magnitude— were required. Second, the scoring accuracy of heterozygote genotypes requires more reads compared to homozygous genotypes (e.g. see Heliconius P3 and Atlantic cod LG01, Figure 2c). Thus, depending on the type of haplotype of the sample, different levels of accuracy are obtained. Third, an increase in scoring accuracy at lower numbers of reads is observed for those comparisons for which more SNPs could be obtained (Table1, Figure 2c). Best scoring accuracy is obtained for the population comparison of Atlantic cod, for which population of origin can be determined with 1000 reads or less than 0.0001x fold nuclear coverage (Figure 2c). Finally, BAMscorer *-score_bams* takes—on average— less than five minutes to complete each comparison.

## Discussion

The BAMscorer program allows genomic assignment on extremely low-coverage sequence data, thereby greatly increasing the capacity for conducting population genomics analysis on poor-quality data. Not only will this expand the amount of information that can be gleaned from such sequences, but it will also reduce waste in the aDNA research pipeline. Sequence data with coverage as low as 0.009x is often discarded as there is little usable information that can be recovered from such poor-quality data. Applying our method will allow these sequences to be used, both reducing waste in the laboratory, and providing a higher degree of confidence that usable information can be recovered when conducting destructive sampling. The method is, additionally, quite fast and can be applied to large quantities of data at one time, providing an efficient overview of the biological characteristics of a large dataset. Such analysis can provide crucial information on population of origin, past trade routes and/or migration patterns, and on species’ ecology and evolution, depending on research question and context.

Additionally, BAMscorer shows promise for application to disciplines outside or adjacent to the field of ancient DNA (e.g. Bohmann et al., 2020). As the underlying methodology is generalistic in design, it is not specific to use with ancient samples. It could therefore be applied to such sequence data as ddRAD (Peterson et al., 2012) or hyRAD (Suchan et al., 2016), two common methods for cost-efficient sequencing used in ecology and evolution studies for modern and historic specimens, respectively. The capacity to quickly identify population of origin, determine between domestic and wild types, assess ecotype distribution, and to identify hybrids could be an incredibly useful tool in the fields of wildlife forensics and conservation genomics.

Our results show that there are several things to take into account when applying the BAMscorer program. Each of our three species showed differentiation in filtering parameters, such as minimum required reads and SNP loading weight cut-off value. We further saw differences in these parameters between cod when looking at the inversion on LG01 as opposed to the genome-wide data. This implies that an understanding of the biological system in question is important for assessing the efficacy of the BAMscorer program. It is further recommended that users explore the filtering parameters as we have done above to ascertain the appropriate parameters for their biological system. The BAMscorer program is further unable to identify de novo inversions and is reliant on existing reference data to create the database from which alignment files are scored.

We have here introduced a novel software program that can be used to increase the information gleaned from extremely low-coverage sequence data. We have found that biological characteristics and genomic assignment can be recovered from sequences with as little as 1000 aligned reads (at ~0.0001x coverage in the case of Atlantic cod). The method is flexible and can be used on various types of genomic data. It is further scalable for BAM files from 1K to 50M reads and can handle up to hundreds of thousands of SNPs without sacrificing computational efficiency. We have shown that it can differentiate between subspecies, ecotypes, and genomic inversions. We expect this approach to be widely applicable in the fields of ancient DNA, conservation genomics and wildlife forensics (Ogden, 2011; Runa & Harbison, 2021).

## Data availability

Reference data for all species have been publicly released earlier and are available from the European Nucleotide Archive (ENA) with the following accession numbers: Heliconius; PRJEB12740 and PRJEB40136, Atlantic herring; PRJNA642736, Atlantic cod; PRJEB29231 and PRJEB41431. The nine ancient Atlantic herring sequences are available at ENA under accession number PRJEB45393.

## Program availability

The full software package is available for download at: https://github.com/laneatmore/BAMscorer

## Acknowledgements

We thank A. Gondek-Wyrozemska for processing the ancient Atlantic herring specimens. We are grateful for the computational resources provided by Saga through allocations to the Centre for Ecological and Evolutionary Synthesis at the University of Oslo. We also thank M. Skage, S. Kollias, M.S. Hansen, and A. Tooming-Klunderud from the Norwegian Sequencing Centre for sequencing and processing of samples. The project benefited from data generated by the RCN-funded project “The Aqua Genome Project” (221734/O30)”. Finally, this project received funding from RCN project “Catching the Past” (262777) and the European Union’s Horizon 2020 research and innovation programme under the Marie Skłodowska-Curie grant agreement No 813383. The European Research Agency is not responsible for any use that may be made of the information it contains.

## Author Contributions

G.F., L.M.A., and BS wrote the manuscript. B.S. conceived of the project. G.F., L.M.A., and B.S. developed the method. L.M.A. wrote the code for the software program. G.F., L.M.A., and B.S. conducted data analysis and visualization. J.H.B. and D.M. provided archaeological material for sequencing. S.J. and K.S.J provided early access genomic sequence data. All authors have read and approved the manuscript.

